# Contrast and pattern adaptation in visual cortex share a common gain control mechanism

**DOI:** 10.1101/2025.11.13.688361

**Authors:** S. Amin Moosavi, Elaine Tring, Dario L. Ringach

**Author notes:** Correspondence: Dario L. Ringach.

## Abstract

Neuronal populations in primary visual cortex adjust their responses to the statistical structure of the environment, including both stimulus contrast and the probability of occurrence of visual patterns. Here we show that, across a wide range of adaptation states, the distribution of population responses is well described by a zero-inflated log-normal model with three parameters: the probability of “silence” *P*_0_, the log-response mean *µ*, and its variance *σ*^2^. Adaptation produces coordinated changes in *P*_0_ and *µ*, whereas *σ*^2^ remains approximately invariant. These coordinated shifts collapse the family of response distributions onto a one-dimensional manifold, consistent with the existence of a common gain mechanism underlying both contrast and pattern adaptation. We further demonstrate that *µ* obeys power-law relationships with stimulus contrast and with orientation probability, and that *P*_0_ varies linearly with *µ*. Finally, we show that these empirical relations arise naturally in a population of linear–nonlinear neurons driven by Gaussian inputs whose mean, but not variance, is modulated by the environment. Together, these results suggest that contrast and pattern adaptation rely on a shared mechanism that adjusts the mean input to cortical populations while preserving the overall structure of their response distribution.

**NEW & NOTEWORTHY:** This study demonstrates that contrast and pattern adaptation in V1 reshape population activity through a common gain mechanism. Despite large changes in responsiveness, the variance of log responses remains invariant, and shifts in mean activity are captured by a simple change in mean input to a population of linear-nonlinear neurons. The proposed mechanism links classic intracellular findings with population-level response distributions.

## INTRODUCTION

We have recently studied how population responses in the mouse primary visual cortex adapt to the probability of occurrence of stimuli, to the distribution of contrasts in the environment, and how these two factors interact [1–3]. These data revealed that the population response magnitude is a separable, power-law of stimulus probability and contrast [2]. In other words, increases in stimulus probability by a certain factor (which tends to reduce the magnitude of the population response) can be offset with increases in stimulus contrast by a different factor (which tends to increase the magnitude), leaving the population magnitude (as measured by its Euclidean norm) invariant. This finding led to the hypothesis that pattern and contrast adaptation may be adjusting one and the same “gain knob” – in other words, tapping into a shared underlying adaptation mechanism. The central goal of the present study is to further explore the existence of such a mechanism and how it could be implemented mechanistically.

To anticipate the results, we report that across all adaptation conditions tested, population response distributions are well-described by a zero-inflated log-normal model characterized by three parameters: *P*_0_, *µ*, and *σ*^2^. Here, *P*_0_ represents the probability of neural “silence” (or no response, given by *P* (*r* = 0)), *µ* is the expected value of the log neuronal response (*µ* = ⟨log(*r*) ⟩ given *r >* 0), and *σ*^2^ is its variance (*σ*^2^ = ⟨log^2^(*r*) ⟩ − *µ*^2^). By measuring how these parameters change between different adaptation states, we found that *σ*^2^ remains approximately invariant, while both *P*_0_ and *µ* co-vary via a power-law relationship. In other words, the family of response distributions across different pattern and contrast adaptation states is effectively one-dimensional, an outcome consistent with the notion of a common mechanism for pattern and contrast adaptation. Finally, we discuss the implications of these findings and introduce a simple linear-nonlinear model with Gaussian inputs consistent with our observations, where the gain “knob” is the mean input to a population of neurons.

## METHODS

All the data analyzed here have been previously published [1–3]. No new animals were used for the present analyses. Here, we provide a brief summary of the experimental designs for completeness. Details about the experiments, including surgical details and data selection, can be found in the original manuscripts.

### Visual Stimulation

In one set of experiments [1], visual stimuli were presented as a sequence of flashed sinusoidal gratings. Each grating was characterized by its orientation, randomly selected from a discrete set of 18 angles spaced evenly from 0^°^ to 170^°^ in 10^°^ increments. Spatial phase was randomly selected from 0^°^ to 315^°^ in 45^°^ increments. The random sequence of gratings was presented at a rate of 3 stimuli per second.

To manipulate stimulus probability distributions across experimental conditions, three different environments were defined in which the probability of each orientation varied. In the “uniform” environment, all orientations were equally likely. In two other environments, specific orientations occurred with higher or lower probabilities, which followed a von Mises distribution, enabling the study of adaptation to environmental statistics in terms of their orientation content.

In a second set of experiments[4], orientations and spatial phases were drawn from a uniform distribution as before, while their contrasts, on the other hand, had different discrete distributions defined on a set of equally spaced values on a logarithmic scale:

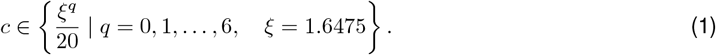

The probability distributions of contrast were truncated log-normal, which varied across three environments (see Fig. 2a, left panels). The geometric mean contrasts of the environments were 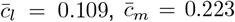, and 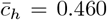, where {*l, m, h* } identify the environments according to their mean contrasts: low, medium and high, defining three environments with different contrast statistics.

In both types of experiments, all six possible permutations of the three environments were presented in random order (18 experimental blocks per session). Each block contained 900 stimuli presented over five minutes (3 stimuli per second), for a total of 5,400 stimuli per environment. A one-minute blank screen was shown between consecutive blocks. Moreover, in both experiments, a spatial frequency of 0.04 cycles per degree was used to match the average preference of V1 neurons [5].

### Data analysis

The response of neuron *i* to a stimulus with orientation *θ* and contrast *c* in trial *t* is denoted as *r*_*i,t*_(*θ, c*). The parameters *P*_0_, *µ, σ*^2^ were estimated by first calculating for each trial as:

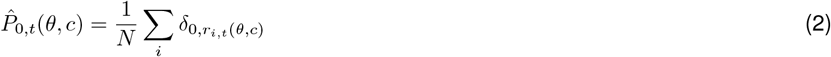

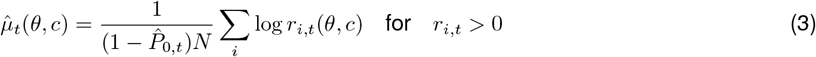

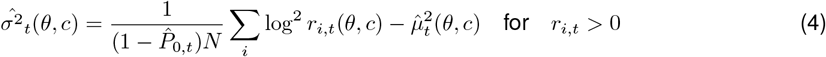

where *N* is the numbers of neurons, and 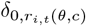 is the Kronecker delta, equal to one if *r*_*i,t*_(*θ, c*) = 0 and zero otherwise. The parameters *µ, σ*^2^, *P*_0_ are then estimated by averaging 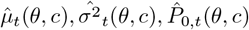 across trials.

To quantify the information conveyed by the population about contrast, we computed the signal-to-noise ratio averaged across orientations:

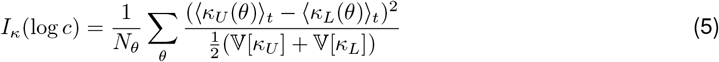

where *N*_*θ*_ is the number of distinct orientation values, *κ* ∈ {*P*_0_, *µ, σ*^2^ } is one of the distribution parameters, and ⟨·⟩ _*t*_ represents an average across trials. The indices *U* and *L* represent two pools of the estimated parameters for contrast values above and below the middle log contrast. Therefore, the information measure represents how informative each parameter is about changing the log contrast values. The variance of one of the estimated parameters across trials is denoted by *V* [*κ*]. Similarly, we computed information about environmental changes:

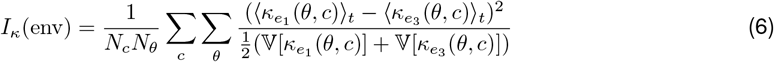

where the indices *e*_1_ and *e*_3_ represent the estimated parameters in the environments with the highest and lowest mean contrast, respectively.

## RESULTS

Our first observation was that the distribution of population responses to any given stimulus and state of adaptation was highly sparse (Fig. 1). Remarkably, in each trial, on average, 80–90 percent of all neurons detected in our field of view showed no response to the stimulus. Second, among the subset of neurons that responded, their activity followed a log-normal-like distribution. This statistical structure was consistent across different environments, stimulus orientations, and contrast values. The approximate log-normality of population responses has been noted in the past and appears to be a widespread feature across different brain areas and species [6]. Our data are consistent with these past observations.

**Figure 1:**
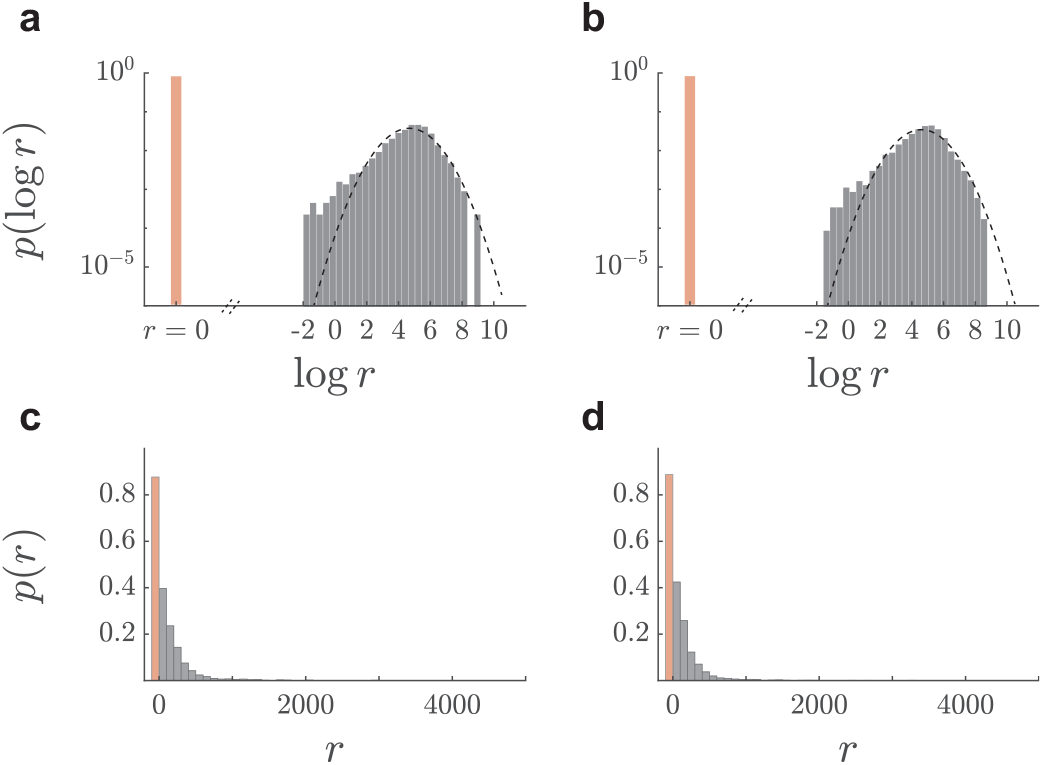
Zero-inflated log-normal distribution of neuronal activity. **a,b** The histograms are evaluated by pooling neuronal activities across trials with the stimulus properties *θ* = 100^°^ and *c* = 0.1357 in one experiment session. In the first row, the orange bar on the left represents *P*_0_, while the nonzero portion is well approximated by a normal distribution, appearing as a parabola when plotted on a logscale *y*-axis. Although there are systematic deviations from log-normality in the skewness of the distribution, a zero-inflated log-normal distribution summarizes our data very well. Panels **a** and **b** are associated with environments with high and medium average contrasts, respectively. Panels **c**,**d** show the same data as in **a**,**b** with linear *x* and *y*-axes, respectively. Note the long tail on the right-hand side of each graph and the sharp peak at zero.

### Adaptation to contrast

The ability to capture the population response distributions using a zero-inflated log-normal distribution allowed us to study how the parameters of the distribution, *P*_0_, *µ*, and *σ*^2^, change when the population was adapted in different environments with different distributions of contrast shown in Fig. 2**a**. In this experiment, orientations were drawn with equal probability and, as a result, the estimated parameters did not show major dependence on this variable (Fig. 2**b,c**). Slight variations across orientations likely reflect the fact that populations did not have a perfectly uniform distribution of preferred orientations. Instead, some orientations are better represented than others. To be able to compare populations with different biases we created datasets where we averaged responses across orientations.

**Figure 2:**
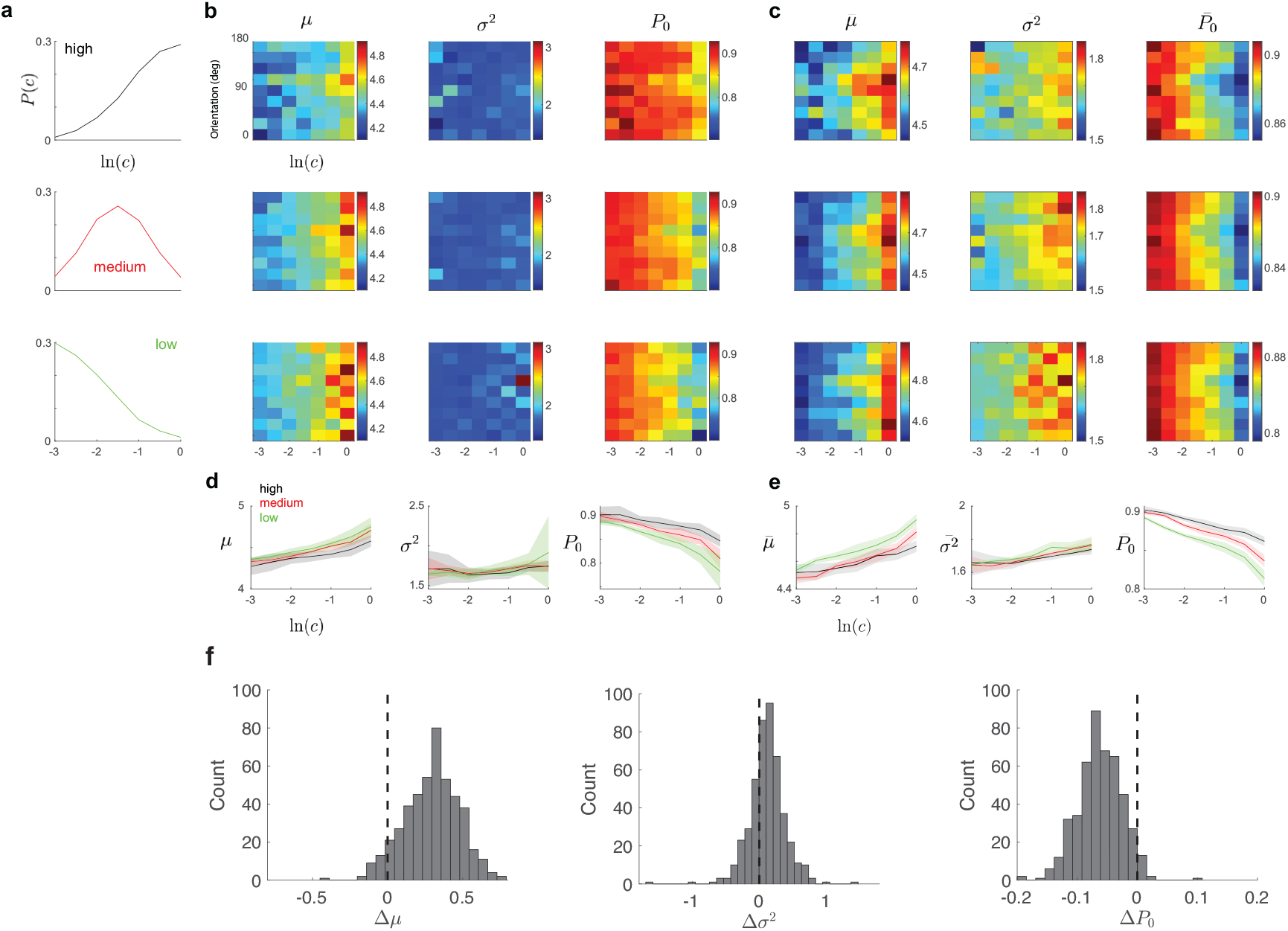
Dependence of zero-inflated, log-normal parameters on orientation and contrast. (**a**) Distributions of contrast in the three environments corresponding to high, medium, and low means. (**b**) Responses in one experiment. The heat maps show dependence of *µ, σ*^2^, and *P*_0_ as a function of log contrast and stimulus orientation. **c** Average dependence of distribution parameters 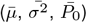 across 17 experimental sessions. (**d**) Mean values of *µ, σ*^2^, and *P*_0_ across orientations as a function of log contrast for the single experiment in panel (**b**). Shaded areas indicate one standard deviation across orientations. (**e**) Mean values of 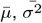 and 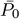 across orientations as a function of log contrast for the population data in panel (**c**). Shaded areas indicate one standard deviation across orientations. (**f**) Histograms show the distribution of parameter variations, Δ*µ*, Δ*σ*^2^, Δ*P*_0_, between high (*c* = 1) to low contrast (*c* = 0.05) across a total of 459 conditions (17 experiments, 3 environments per experiment, and 9 orientations per environment).

A typical example and the average behavior of parameters with contrast increase and the average behavior across all experiment sessions are shown in Figure. 2**d,e**. We see a systematic trends in *µ, P*_0_ with contrast, and to a lesser extent in *σ*^2^ when we focus on the average population data. To examine these changes in more detail, we plotted histogram of the difference in parameters between high and low contrasts (Figure 2**f**). We found that *µ* and *P*_0_ have a strong dependence on contrast with an increase of Δ*µ* = 0.294 ± 0.181 (mean ± SD, Wilcoxon signed-rank test, *n* = 459, *p* − val = 1.19 × 10^−72^), and a decrease of Δ*P*_0_ = 0.062 ± 0.036 (mean ± SD, Wilcoxon signed-rank test, *n* = 459, *p* − val = 5.1 × 10^−75^). On the other hand, *σ*^2^ was more stable, exhibiting a statistically significant, but less pronounced increase, Δ*σ*^2^ = 0.12 ± 0.27 (mean ± SD, Wilcoxon signed-rank test, *n* = 459, *p* − val = 2.9 × 10^−22^). The effect sizes, defined by the mean shift divided by the standard deviation are *>* 3.5 times larger for Δ*µ* (Cohen’s *d* = 1.61) and Δ*P*_0_ (Cohen’s *d* = 1.72), compared to the increase experienced by Δ*σ*^2^ (Cohen’s *d* = 0.44).

Next, we computed an information measure that captures how well we can discriminate between different contrast levels given the parameters of the distribution (see **Methods**). Across experimental sessions, *P*_0_ and *µ* conveyed most of the information about contrast discriminability, whereas *σ*^2^ conveyed very little (Fig 3a-c). Moreover, the same two parameters (*P*_0_ and *µ*) also conveyed information about the environment, thus encoding the adaptation state of the cortical population (Fig. 3d).

**Figure 3:**
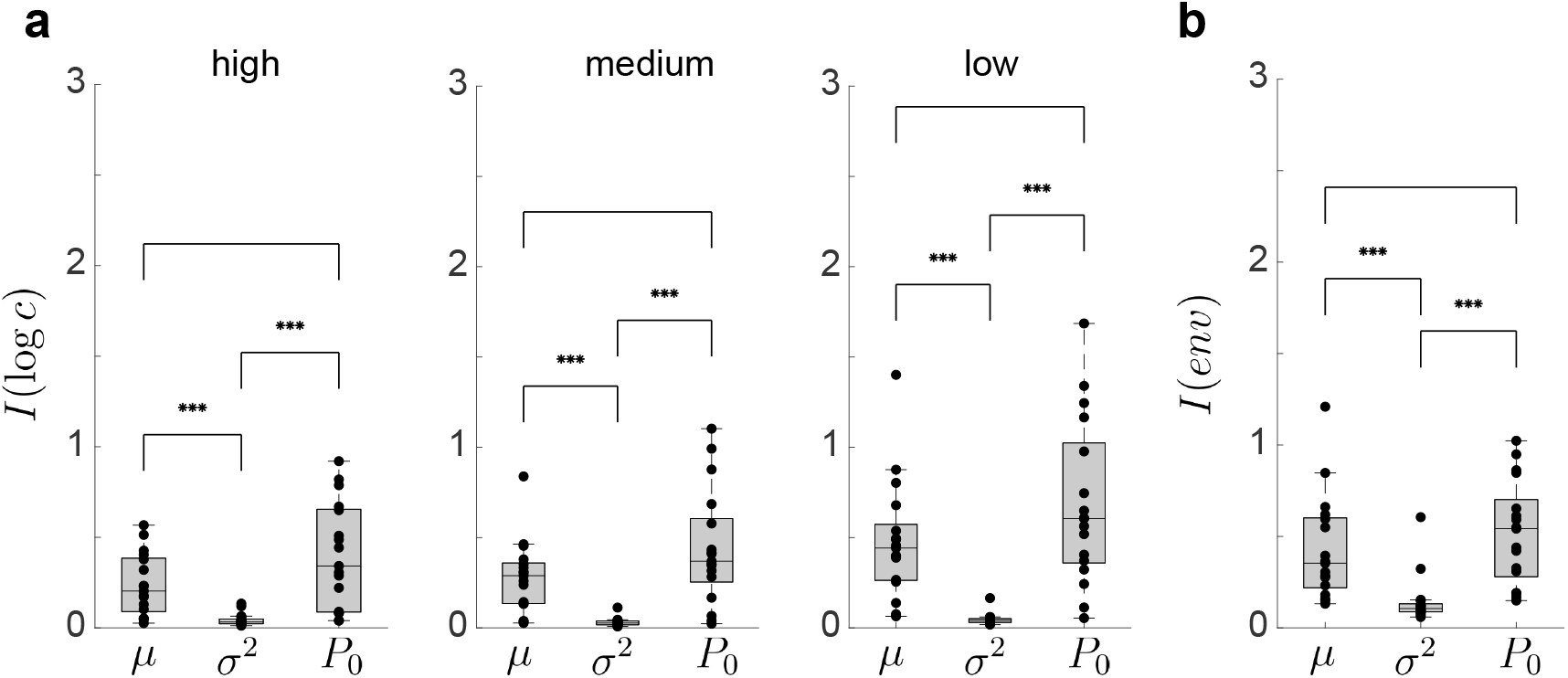
Information about contrast variations and gain modulation is conveyed primarily by *P*_0_ and *µ*, while *σ*^2^ provides negligible information. (**a-c**) Information about log *c* conveyed by *µ, σ*^2^, and *P*_0_ in environments with high, medium, and low prevailing contrasts. (**b**) Information about the environment conveyed by each parameter. In each box-plot, the center line indicates the median, while the bottom and top edges of the box represent the 25th and 75th percentiles. Points beyond the whiskers represent outliers. Stars indicate confidence levels of two-sample Kolmogorov-Smirnov tests. Three stars indicate p-value of *<* 0.001, indicating a significant difference between the group of *σ*^2^ and either *µ* or *P*_0_. No star means the null hypothesis cannot be rejected.

The relationship between *µ* and contrast was well captured by a power-law function (Fig. 4**a,b**):

**Figure 4:**
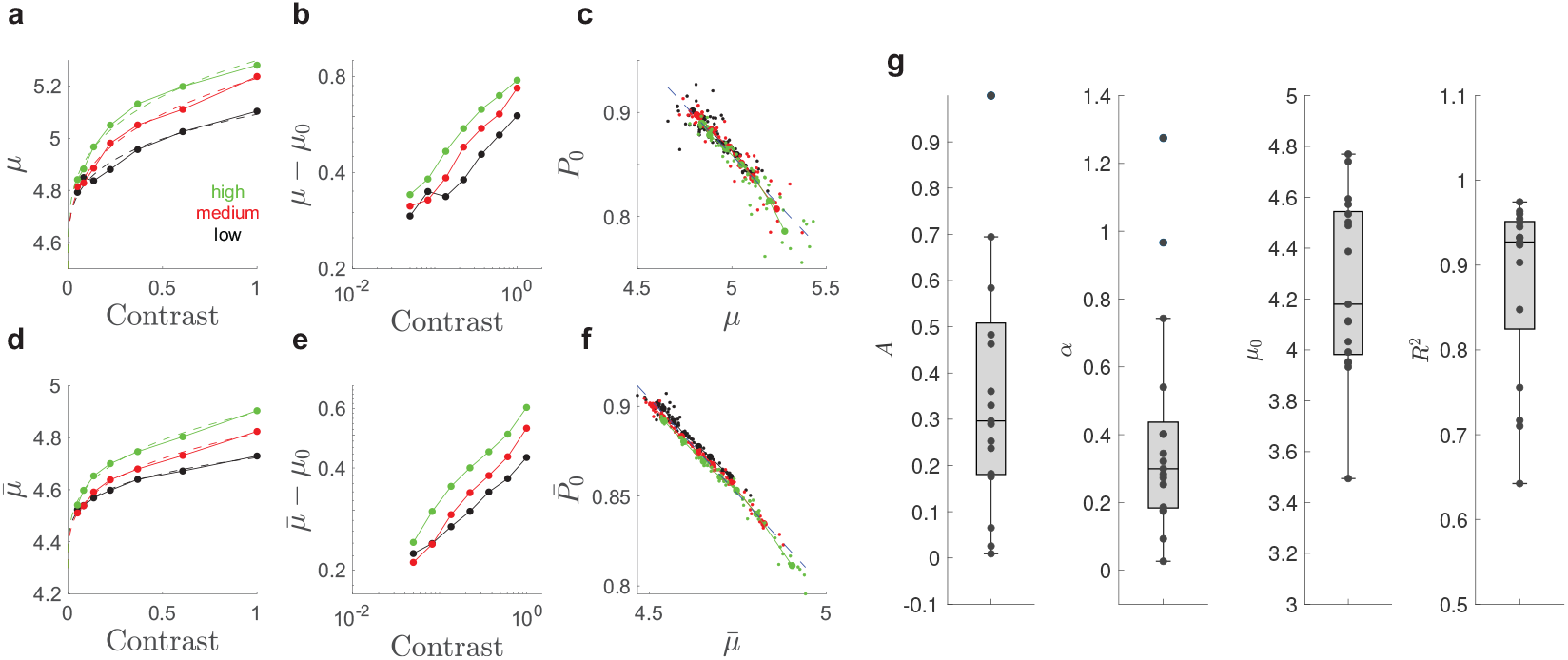
Power-law model captures the behavior of *P*_0_ and *µ*. (**a-c**) Results from a representative experiment session (one of 17). (**a**) *µ* averaged across orientations and plotted versus contrast for three environments (colors correspond to different environments). Dashed lines show fitted power-law functions. (**b**) The same data as **a** is used to plot *µ* − *µ*_0_ versus contrast on a log-log scale. (**c**) Linear relationship between *P*_0_ and *µ*. Each dot represents estimated parameters for a given orientation and contrast. Larger circles denote average values across orientations. The dashed line is a linear fit to all the points. Colors represent different environments and match the legend in (**a**). (**d-f**) Same as (**a-c**), but averaged across all 17 experimental sessions, with parameters denoted as 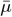 and 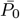 **(g)** Fitted parameters of the power-law function (Equation 7), *A, α, µ*_0_, and the *R*^2^ as a goodness of fit measure are shown from left to right, respectively. In each boxplot, the center line represents the median. The bottom and top edges of the box represent the 25th and 75th percentiles, respectively. Points beyond the whiskers represent outliers.

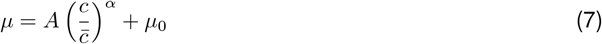

where *A, α*, and *µ*_0_ were simultaneously fitted to the data of all environments in each session (Figure 4 **a**). The fitted values and variations across all 17 experiment sessions are shown in (Figure 4 **g**). We find the median values for parameters across all experiment sessions as *med*(*A*) = 0.30, *med*(*α*) = 0.30, and *med*(*µ*_0_) = 4.17. The *R*^2^ of the fit is close to one with a median of 0.93, indicating an excellent fit in most experiment sessions (Figure 4 **g** right panel). As a result, the log-log plots of *µ* − *µ*_0_ versus contrast in different environments are linear with similar slopes (Figure 4 **b**). Additionally, we observed a strong linear relationship with a negative slope between *P*_0_ and *µ* (Figure 4 **c**), indicating an increase in the number of responding neurons with an increase of contrast or a decrease of the geometric mean contrast of the environment, 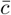. Furthermore, we confirmed the overall validity of the above power-law relation, Equation 7, by fitting it to 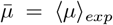 versus contrast (Figure 4 **d-e**). Brackets indicate the average of *µ* across all experiment sessions. Similarly, the overall average linear relation between 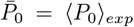 and 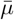 is also confirmed in Figure 4 **f**. In summary, these findings indicate that contrast gain modulation can be understood as a transformation in which the geometric mean of the contrast 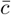 of an environment rescales contrast values [4]. Such a transformation can be interpreted as a translation of the log-normal component of the distribution along the log-response axis, accompanied by a corresponding linear change in *P*_0_, while maintaining the overall shape of the distribution.

### Adaptation to orientation

Neuronal populations in primary visual cortex also adapt to the probability of occurrence of a stimulus in an environment. In the case of oriented, sinusoidal gratings, we previously showed that a power law links the magnitude of the population response and probability of a stimulus [1]. With this result in mind, we examined how the parameters *µ, σ*^2^, *P*_0_ would change across environments defined by different probability distributions in the orientation domain. Will we observe a correlated change in *µ* and *P*_0_ and an invariant *σ* as during adaptation to contrast, or will the responses show a different pattern? To answer this question, we analyzed experiments where the probability distributions in two environments, A and B, defined by von Mises with the same variance, peaking at 0^°^ and 90^°^, respectively, while the distribution in a third environment C was uniform (Figure 5**a**). In these experiments, the contrast was fixed at 100%.

**Figure 5:**
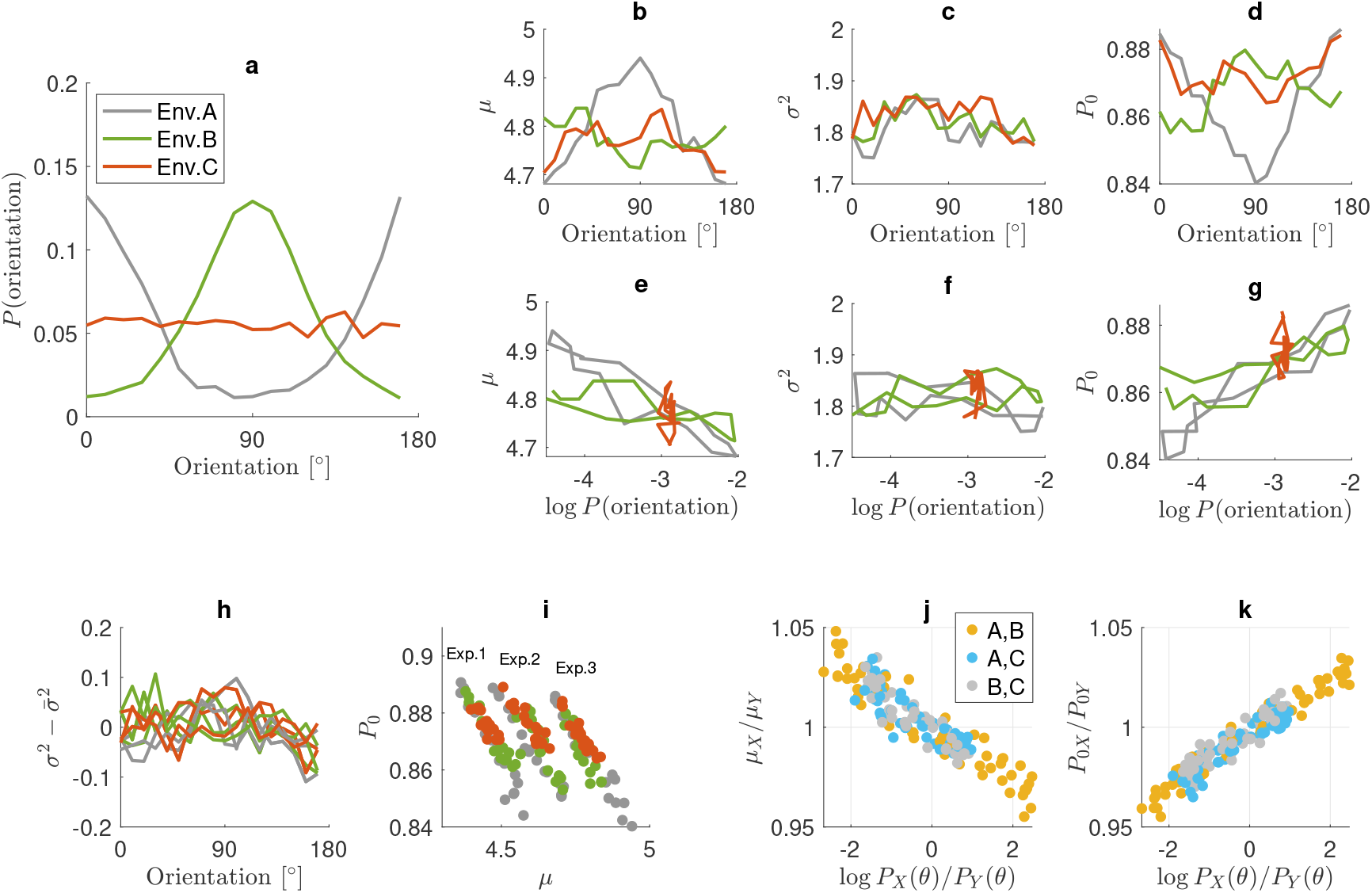
Adaptation to stimulus orientation in three environments. **A, B, C**. Colors gray, red, and green are consistently used in all panels of this figure. (**a**) Probability distribution of orientations in the three environments composed of gratings with orientations in {0, 10, 20, …, 170 } degrees. (**b-d**) Variations of the parameters *µ, σ*^2^, *P*_0_ with respect to orientation for the three environments. (**e-g**) Relation between the three parameters *µ, σ*^2^, *P*_0_ and log probability of orientations. (**h**) Variations of *σ*^2^ around its mean 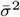 for three experiments with the same environment setup as in **a**. The result suggests invariance with respect to orientations and environments. (**i**) Linear relation between *P*_0_ and *µ* is preserved in all experiments with the same slope. (**j**) The ratio between *µ* parameters in different pairs of environments *X, Y* ∈ {*A, B, C* } versus the ratio of the probability of orientations. Colors orange, light blue, and light gray represent the ratios between environments (A,B), (A,C), and (B,C), respectively. (**k**) The ratio between *P*_0_ parameters in different pairs of environments versus the ratio of the probability of orientations. Same coloring scheme as in **j** is used.

We found that, depending on the probability of the stimuli, *µ* and *P*_0_ systematically varied with orientation (Figure 5**b,e,d,g**) and, as before, *σ*^2^ did not show any systematic change (Figure 5**c,f**). This was consistently observed in all experiment sessions (Figure 5**h**). Specifically, *µ* and *P*_0_ turned out to have negative and positive correlations with the probability of orientations, respectively (compare Figure 5**d,g** with Figure 5**b,e**). Interestingly, similar to the experiments with contrast adaptation, discussed above, there was a linear relation between *µ* and *P*_0_ that effectively constrains the changes in parameter space of (*µ, σ*^2^, *P*_0_) to a one-dimensional manifold (Figure 5**i**).

Although the environment C was defined by a uniform distribution in the orientation domain, some experiments showed systematic variations of *µ* and *P*_0_ with orientation. For example, the red plot in Figure 5**b,d** suggests that the responses become stronger around horizontal (90^°^) orientation (larger *µ* and smaller *P*_0_). Such bias arises from a higher proportion of neurons having a preferred orientation tuned to horizontal compared to other orientations [7–10]. To examine the data in a way that factors out such biases we plotted the ratio of the parameters (*µ*_*X*_ */µ*_*Y*_ and *P*_0*X*_ */P*_0*Y*_) as a function of the log of the ratio of the orientation probabilities *P*_*X*_ (*θ*)*/P*_*Y*_ (*θ*) across pairs of environments with *X, Y* ∈ {A,B,C } (Figure 5**j,k**). The data of all ratios collapsed on one line; hence, they were independent of the bias factor, which must be multiplicative and canceled out in the ratios.

### A model for emergence and adaptation of zero-inflated log-normal distributions

One may wonder, what kind of mechanistic model may give rise to a zero-inflated log-normal distribution? Following the approach in [11], we introduce a simple linear-nonlinear model that exhibits zero-inflated lognormal response distributions and captures the observed behavior of *P*_0_, *µ, σ*^2^ in the data. The theory suggests that the distribution of input to the neuronal populations in V1 undergoes distinct changes in response to contrast and environmental variations that control the distribution of cortical firing rates.

We considered a population of *N* neurons in the V1 area receiving external stimulus-driven inputs as well as recurrent local inputs from other neurons. In each trial *t*, each neuron *i* receives the input *I*_*i,t*_. We assumed that the input distribution across neurons was normal, i.e., *I*_*i,t*_ values were sampled from a normal distribution with mean *Ī* and variance Σ^2^. In addition, we considered a non-linear thresholded input-response relation for all neurons in all trials:

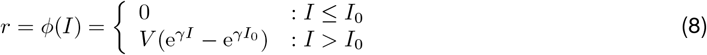

parametrized by *V, γ*, and *I*_0_. In the above equation, the term 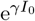 was set to make the function continuous. Using the normal distribution function of *I*

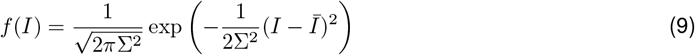

and *ϕ*(*I*), it is straightforward to calculate the probability density function of *r* as

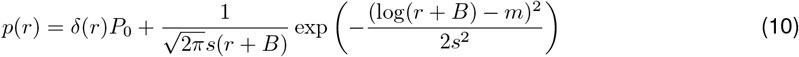

in which *δ*(·) is the Dirac delta function accounting for the zero-inflated component (See Figure 6**a,b**). The non-zero component is a shifted (three-parameter) log-normal distribution. We note that the log-normal section of the mixture is truncated because *r >* 0. In addition, normalization of *p*(*r*) dictates 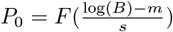 where *F* (·) is cumulative standard Gaussian distribution. Therefore, there are only three free parameters *P*_0_, *m, s* characterizing the distribution function. The relations between the parameters of the distribution and the model are summarized as

**Figure 6:**
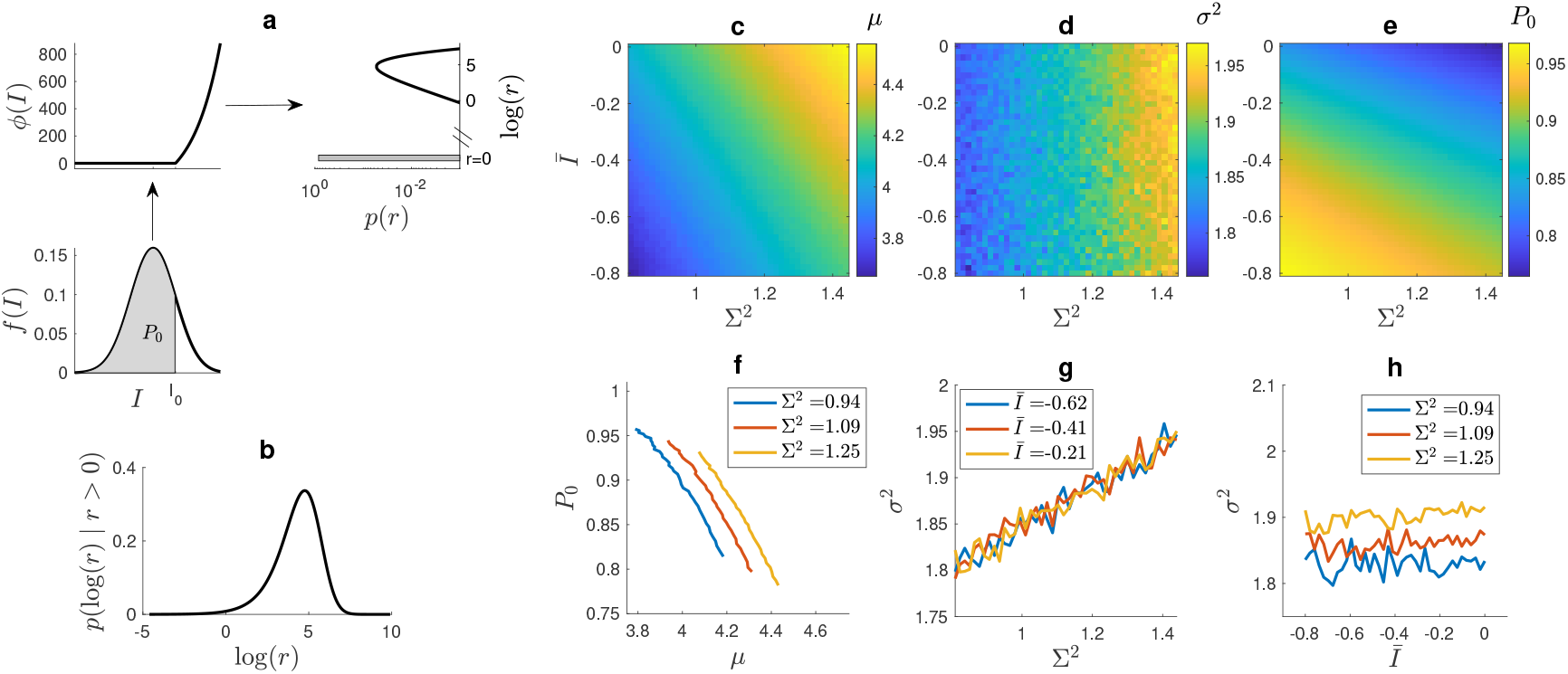
Gaussian input nonlinear output model. (**a**) Gaussian input *f* (*I*) together with the non-linear input-response function *ϕ*(*I*) lead to a zero-inflated log-normal-like response distribution. (**b**) Conditional probability density function *p*(log(*r*) | *r >* 0) of the responses (Equation 14) **a**. (**c,d,e**) Behavior of *µ* (**c**), *σ*^2^ (**d**), and *P*_0_ (**e**) versus *Ī* and Σ^2^. The range of *Ī* and Σ^2^ is chosen such that *µ, σ*^2^, *P*_0_ cover the range observed in population responses of V1 neurons. (**f**) Approximately linear behavior of *P*_0_ and *µ* for three fixed values of Σ^2^. (**g**) The linear relation of *σ*^2^ and Σ^2^ is independent of *Ī*.(**h**) For fixed Σ^2^, the parameter *σ*^2^ is independent of *Ī*.

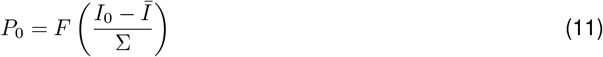

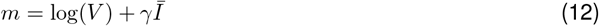

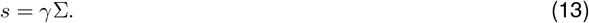

Based on Eq 10 we write the conditional probability density function (See Figure 6**b**)

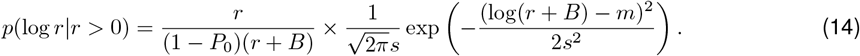

The parameters *µ* and *σ*^2^ are written as expectation values:

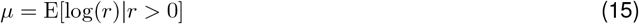

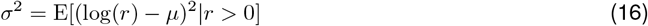

that give the relation between (*µ, σ*^2^) and the parameters (*m, s*^2^, *P*_0_), and consequently the parameters of the model (*V, I*_0_, *γ, Ī*, Σ). Keeping the parameters of the input-response function (*V, I*_0_, *γ*) fixed, we studied how *µ, σ*^2^, and *P*_0_ vary by changing the input mean *Ī* and variance Σ^2^. We found that *µ* and *P*_0_ depend on both *Ī* and Σ^2^ (Figure 6**c,e**), and there is an approximately linear relationship between *µ* and *P*_0_ (Figure 6**f**). However, *σ*^2^ only depends on Σ^2^ linearly and does not show any variations with respect to *Ī* (Figure 6**d,g,h**).

Then, we focused on setting the model parameters to capture the observed adaptive behaviors in the neuronal populations. We considered three key properties in the population response adaptations: (1) *σ*^2^ is invariant across environments, orientations, and contrast values. (2) There is a one-to-one, approximately linear, relation between *P*_0_ and *µ*. While the slope of the line is invariant across environments, the range of *P*_0_ and *µ* varies by changing environments (changing with average contrast (Figure 2**d,e**) and probability of orientations (Figure 5**i**). (3) *µ* and *P*_0_ are power-law functions of 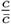 and 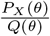 with *Q*(*θ*) being a symmetric concave function with a maximum at *θ* = 90^°^, accounting for the horizontal orientation bias. Interestingly, as shown in Figure 6**f-h** points (1) and (2) are already built into the model. We can keep *σ*^2^ invariant by fixing Σ^2^. In other words, the variance of the input distribution must be kept fixed in all conditions, regardless of the environment, orientation, and contrast. To further study the adaptations, we formulated *Ī* as the only parameter that varies with environment, contrast, and orientation to capture the observed power-law behavior of *µ*. Therefore, because *µ* is an increasing function of *Ī* at fixed Σ^2^ (Figure 6**c**) we chose the following formula

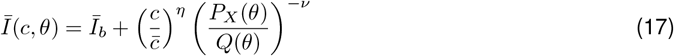

where *Ī* _*b*_ is the baseline mean input that is achieved at *c* = 0, i.e. absence of stimulus. The exponents *η, ν* are both positive, accounting for proportional and inverse relation with contrast and orientation probability, respectively (See Figure 4**a,d** and 5**e,g**). In addition, we chose the orientation bias function

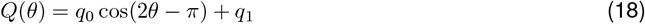

which is symmetric and peaks at *θ* = 90^°^. To avoid negative or zero values, we set *q*_1_ *>* 2*q*_0_ (*q*_0_ = 0.9 and *q*_1_ = 1.9).

The model captures the changes in gain observed in the experiments (Figures 2,4,Figure 7). The environments simulated were the same as those used in the experiments. A power-law relation between *µ* and 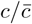 was verified by fitting the power-law function of Equation 7 to the simulated data (Compare Figure 4**a** and Figure 7**b**). The value of *σ*^2^ remained invariant across environments and contrast values (Figure 7**c**). As shown in Figure 7**d**, *P*_0_ was a decreasing function of contrast, with the curves having the same trend as shown in Figure 2**d,g**. Estimated *µ* and *P*_0_ fell on a one-dimensional manifold in the parameter space of (*µ, σ*^2^, *P*_0_) (Figure 7**e**).

**Figure 7:**
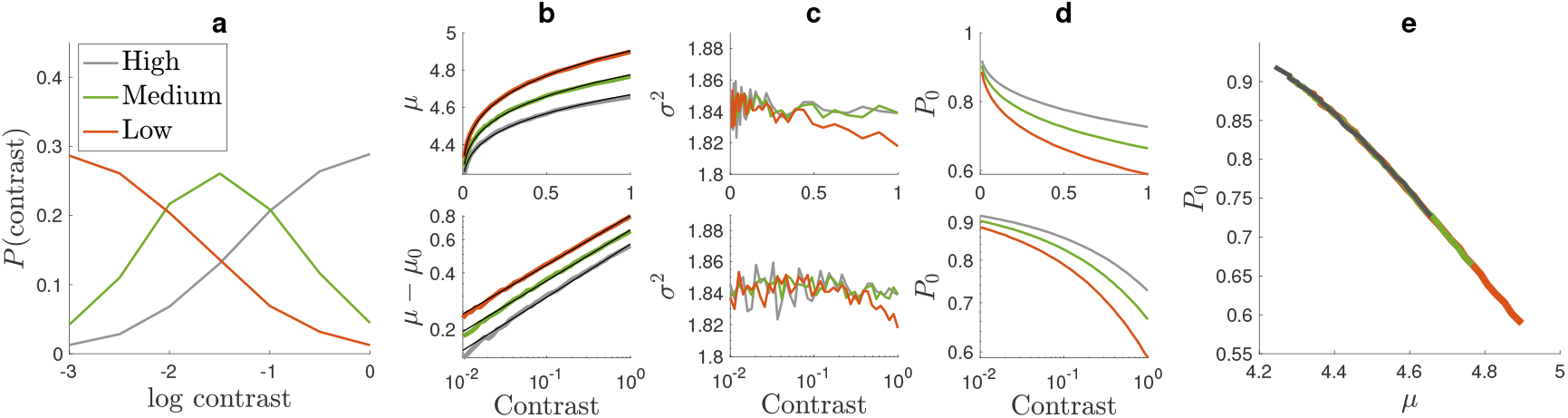
Modeling adaptation to contrast. The same three environments are used as in the experiments shown in Figure 2. The three environments with low, medium, and high contrasts are shown respectively in green, red, and gray. Stimulus orientations are equally distributed. (**a**) Probability distribution of contrast in the three environments. (**b**) Behavior of *µ* as a function of contrast in the three environments. The top and bottom panels show plots in linear and log scale, respectively. Solid black lines are power-law fits according to Equation 7 with *α* = 0.36 and *µ*_0_ = 4.1. (**c**) Same convention as in **b** but for *σ*^2^, which remains invariant across environments and contrast values. (**d**) Decreasing behavior of *P*_0_ with contrast. Same conventions as in **b,c**. (**e**) Linear, one-to-one relation between *P*_0_ and *µ*. Note that the range of both parameters increases by decreasing 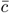 from high-to low-contrast environments. In all panels to replicate the average range of *µ, σ*^2^, *P*_0_ observed in brain recordings, we set *γ* = 0.8, *I*_0_ = 1.5, and *A* = 100. To generate this figure, we keep Σ = 1 and let *Ī* vary according to equation 17 with *η* = 0.15.

The model also captures the adaptation to stimulus orientations. We considered the same environments used in experiments with probabilities plotted in Figure 8**a**. Gray, red, and green indicate environments A, B, and C, respectively. Behavior of *µ, σ*^2^, and *P*_0_ shown in Figure 8**b,c,d** closely followed the experimentally observed behavior shown in Figure 5**b,c,d**. Similarly, the relation of *µ* and log *P* (*θ*) as well as that of the ratios of parameters were well captured by the model (Figure 8**e-g**). Furthermore, our model also gave the power-law relation between the ratios of the population norms *r*_*X*_ */r*_*Y*_ and the respective probabilities for different environment pairs in agreement with our previous results in [1] (Figure 8**h**).

**Figure 8:**
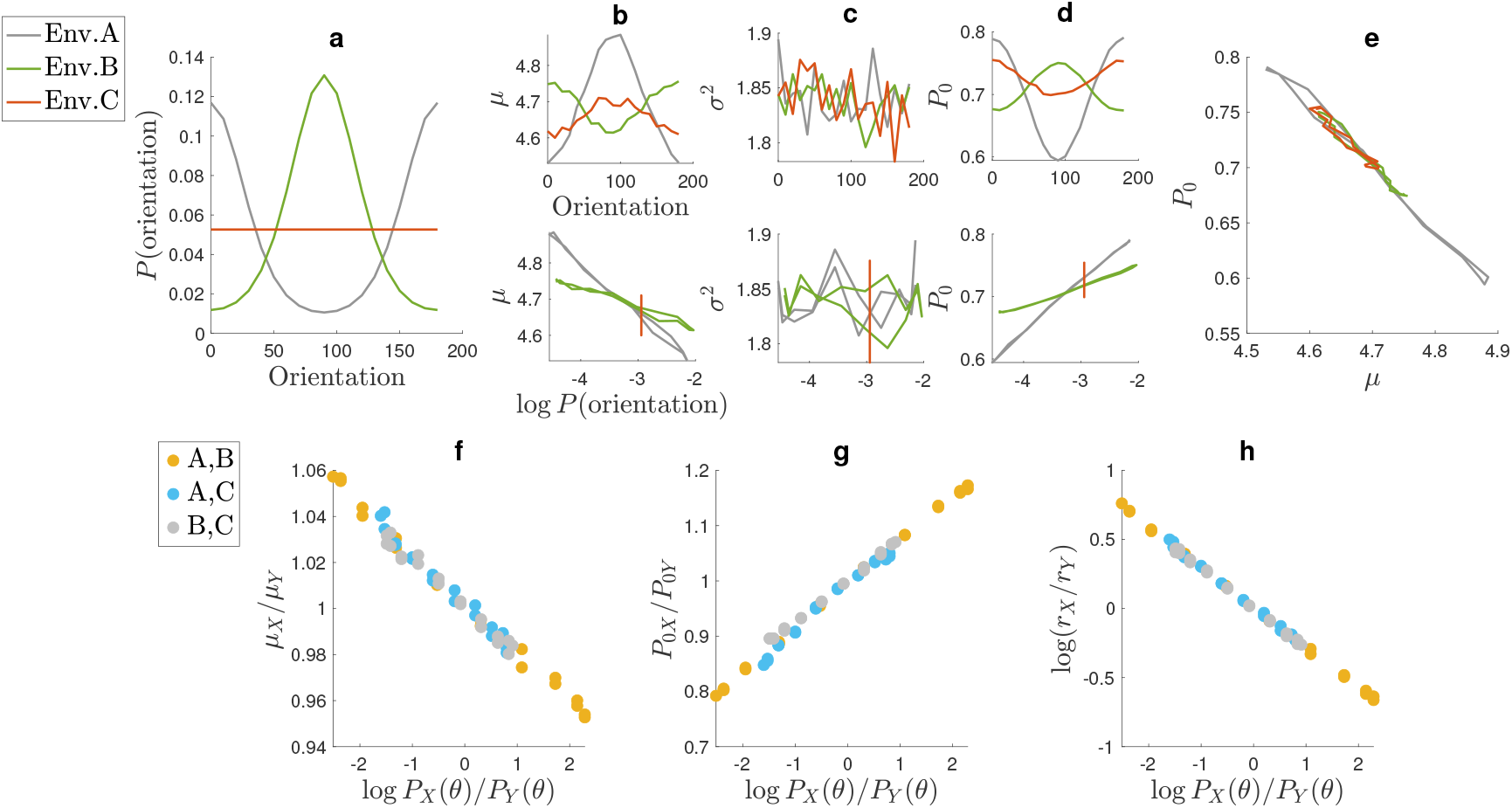
Modeling the adaptation to orientation. The three environments A, B, and C are shown respectively by gray, red, and green as in Figure 5. In these environments, the contrast value is equal to 1. (**a**) Probability distribution of orientations in the three environments. (**b-d**) Behavior of *µ, σ*^2^, and *P*_0_ versus orientation (top panel) and orientation log probability (bottom). Note the orientation bias in the homogeneous environment C, shown in red. (**e**) Linear relation of *µ* and log probability of orientation. Regardless of the environment, the parameters fall on a one-dimensional manifold in the *P*_0_ and *µ* space. (**f,g**) The ratios between the parameters evaluated in different environments versus the ratios of the log probability of orientations are linear for both *µ* (**f**) and *P*_0_ (**g**) parameters. **(h)** Power-law relation between the ratios of mean neuronal response and ratios of probabilities as reported in [1]. In all panels to replicate the average range of *µ, σ*^2^, *P*_0_ observed in brain recordings, we set *γ* = 0.8, *I*_0_ = 1.5, and *A* = 100. We keep Σ = 1 and let *Ī* vary according to equation 17 with *ν* = 0.1.

Overall, as the model captures all salient aspects of neuronal adaptations of the population response distributions in the V1 area, it can be considered a strong candidate for explaining the underlying mechanisms of adaptations. The model suggests that the sparseness and long-tail distribution of neuronal responses are the result of a non-linear input-output function of neurons, such that in a given trial, most neurons remain silent while receiving subthreshold inputs (*I < I*_0_). In addition, within this setting, the invariance of *σ*^2^ is due to the invariant width of the Gaussian input distribution. In other words, across adaptation states, we predict a constant input variance across all conditions. This leaves the mean input current *Ī* as the only parameter to be used by the brain in contrast gain control and orientation adaptation that acts as one control knob to adapt to different environments, modeled by Equation 17.

## DISCUSSION

Adaptation in single V1 neurons has been studied in detail uncovering how tuning functions change in response to pattern adaptation [12–34] and how the contrast response curves shift the mean contrast of the environment is varied [12, 35–39]. In these classic studies, stimulus parameters were optimized to maximally drive individual neurons, and the data did not fully capture how adaptation influences the distribution of responses at the population level. Given the sharp selectivity of neurons to stimulus parameters, one naturally expects the response to any one stimulus to evoke a strong response in a small fraction of cells, while the vast majority would respond weakly or not at all. Our recent work has focused on how pattern and contrast adaptation affect the magnitude (the Euclidean norm) and direction of the mean population response vector. Our findings revealed that contrast gain can be well described as a re-parametrization of the contrast response function of neuronal populations [4], and that pattern adaptation reflects the probability of a stimulus via a power-law function [1, 3].

The goal of the present study was to analyze how pattern and contrast adaptation affect the distribution of responses in V1 populations. We found that this distribution is well-described by a zero-inflated log-normal function characterized by three parameters *P*_0_, *µ, σ*^2^. The parameter *σ*^2^, is largely invariant under adaptation. On the other hand, *µ* and *P*_0_ change within and across environments. These changes follow power-law functions in which the changes in the environment contrast are equivalent to rescaling the contrast values by their geometric mean, and changes in probabilities are rescaled by a “default” orientation preference function, reflecting inherent biases in the population. Such re-parametrization of population response statistics agrees with our previous work on population response vectors [1, 4].

Our analyses also align with past intracellular recordings from cat V1, which have shown that contrast adaptation shifts in the neuronal operating such that prolonged exposure to high-contrast stimuli produces a sustained hyper-polarization of the membrane potential and a reduction in mean synaptic drive, while the variance of membrane fluctuations remains largely unchanged [40, 41]. This hyper-polarization translates the neuron’s input–output function along the voltage axis, producing a rightward shift of the contrast–response curve without altering its slope—consistent with divisive normalization; a behavior that could be replicated by a rectification model. Here, we considered an analogous model working at the population level, which shows how a zero-inflated log-normal-like distribution can emerge in neuronal populations of linear-nonlinear units that receive normally distributed inputs [11]. We further demonstrated how adaptation can be explained by considering only the orientation and contrast dependence of the mean *Ī* of this normal input. Altogether, the data are consistent with the notion that pattern and contrast adaptation rely on the same underlying mechanism, where contrast (*c*) and orientation probability *P* (*θ*) play similar roles in setting the operating point of neurons by modulating the mean input to the population.

Sparse population responses with heavy-tailed activity distributions have long been proposed as a hallmark of efficient coding—minimizing metabolic cost while preserving representational fidelity and dynamic range [42–46]. Indeed, in recent work, we showed that such efficiency principles naturally give rise to the empirically observed power-law relationship between stimulus probability and population response magnitude in mouse V1 [47] (see also [48] for another application of efficient coding to adaptation). The mechanistic model presented here reaches the same conclusion from the opposite direction: by linking adaptation to changes in the mean input current of a fixed-variance Gaussian drive, it reproduces the same universal power-law scaling observed experimentally. That two independent approaches—a top-down, normative theory and a bottom-up, biophysical model—converge on the same principle suggests a deep correspondence between the nonlinear input–output dynamics of cortical neurons and the global objective of energy-efficient coding. Together, these results point to a unifying framework in which contrast and pattern adaptation emerge as expressions of the same underlying law governing efficient population responses in primary visual cortex.

Before concluding, it is important to acknowledge some limitations of the present study. First, our analyses are based on two-photon calcium imaging, which provides an indirect measure of neuronal activity. The recorded fluorescence signals reflect intracellular calcium transients rather than action potentials. As a result, the inferred distribution of population responses, including the zero-inflated log-normal fits, should be interpreted in that context. In particular, the nonlinear relationship between spike rate and fluorescence amplitude, together with potential thresholding effects of the spike-inference algorithm for weak signals, may distort the exact form of the response distributions. Future electrophysiological recordings would be needed to confirm that the same relationships hold at the level of spike counts. Second, the adaptation paradigms used here involved continuous presentation of diverse stimuli, with contrasts and orientations randomly interleaved within each block. This design probes adaptation to the statistical structure of the environment rather than to a single stimulus presented for an extended period of time. Thus, the observed effects likely reflect rapid, ongoing gain adjustments across the population, rather than the slower, steady-state forms of adaptation typically observed after prolonged exposure to a fixed, visual pattern. While our approach offers a closer approximation to natural visual statistics, it does not capture all timescales or mechanisms of adaptation. Finally, our analyses aggregate responses across hundreds of neurons within each field of view of optical sections within layers 2/3. Future work combining cell-type–specific labeling or depth-resolved imaging with electrophysiological validation could help determine how the relationships differ between cell types and cortical layers.

## Data Availability

Data have already been published in a figshare repository. The code will be published as well, following acceptance of the manuscript.

## Author Contributions

D.L.R. and S.A.M. conceived and designed the study; E.T. and D.L.R. performed the experiments; D.L.R. and S.A.M. did the modeling and analyzed the data; D.L.R. and S.A.M. drafted the manuscript; All authors approved the final version of the manuscript.

